# Distance Indexing and Seed Clustering in Sequence Graphs

**DOI:** 10.1101/2019.12.20.884924

**Authors:** Xian Chang, Jordan Eizenga, Adam M. Novak, Jouni Sirén, Benedict Paten

## Abstract

Graph representations of genomes are capable of expressing more genetic variation and can therefore better represent a population than standard linear genomes. However, due to the greater complexity of genome graphs relative to linear genomes, some functions that are trivial on linear genomes become more difficult in genome graphs. Calculating distance is one such function that is simple in a linear genome but much more complicated in a graph context. In read mapping algorithms, distance calculations are commonly used in a clustering step to determine if seed alignments could belong to the same mapping. Clustering algorithms are a bottleneck for some mapping algorithms due to the cost of repeated distance calculations. We have developed an algorithm for quickly calculating the minimum distance between positions on a sequence graph using a minimum distance index. We have also developed an algorithm that uses the distance index to cluster seeds on a graph. We demonstrate that our implementations of these algorithms are efficient and practical to use for mapping algorithms.

## 1 Introduction

Conventional reference genomes represent genomes as a string or collection of strings. Accordingly, these so-called “linear reference genomes” can only store one allele at each locus. The resulting lack of diversity introduces a systematic bias that makes samples look more like the reference genome [20]. This reference bias can be reduced by using pangenomic models, which incorporate the genomic content of populations of individuals [1]. Sequence graphs are a popular representation of pangenomes that can express all of the variation in a pangenome [13]. Sequence graphs have a more complex structure and contain more data than linear genomes. This tends to make functions on a sequence graph more computationally challenging than analogous functions on linear genomes.

One such function is computing distance. In a linear genome, the exact distance between two loci can be found by simply subtracting the offset of one locus from the offset of the other. In a graph, calculating distance is much more complicated; there may be multiple paths that connect the two positions and different paths may be relevant for different problems.

Distance is a basic function that is necessary for many functions on genome graphs; in particular, calculating distance is essential for efficient mapping algorithms. In a seed-and-extend paradigm, matches between the query sequence and reference are used to identify small regions for expensive alignment algorithms to align to [17, 10, 15, 6, 18, 16]. Often these regions are identified by clusters of matches. Clustering requires repeated distance calculations between seeds and can be very slow in graphs as large as whole genome graphs. The prohibitive run time of clustering algorithms can make them impractical for mapping and some mapping algorithms omit this step entirely [16].

We have developed an algorithm to calculate the minimum distance between any two positions in a sequence graph and designed a index to support it. We also developed a clustering algorithm that clusters seeds based on the minimum distance between them. Our algorithms are implemented as part of `vg`, a variation graph toolkit [6].

## 2 Background

### 2.1 Sequence Graph Structure

A sequence graph is a bidirected graph in which each node is labeled by a sequence of nucleotides. A node *X* has two sides, 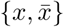. For convenience, we will consider *x* to be the “left” side and 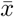 to be the “right”. This induces a directionality on *X*, so that we may consider a left-to-right traversal of *X* to be forward, and a right-to-left traversal backward. However, we note that the designation of “left” and “right” is arbitrary. They can be swapped without changing the underlying graph formalism. Conceptually, a forward traversal corresponds to the forward strand, and a backward traversal corresponds to the reverse strand.

Paths in a bidirected graph must obey restrictions on both nodes and edges. Edges connect two node sides rather than nodes. A path consists of an alternating series of oriented nodes and edges. The path must enter and exit each (non-terminal) node through opposite node sides. In addition, there must exist an edge connecting each node side that is exited with the next node side that is entered. In Figure 1, the graph has an edge between *ā* and *c*. A path including this edge would go from *A* to *C* traversing both forward, or from *C* to *A* traversing both backward.

**Fig. 1.**
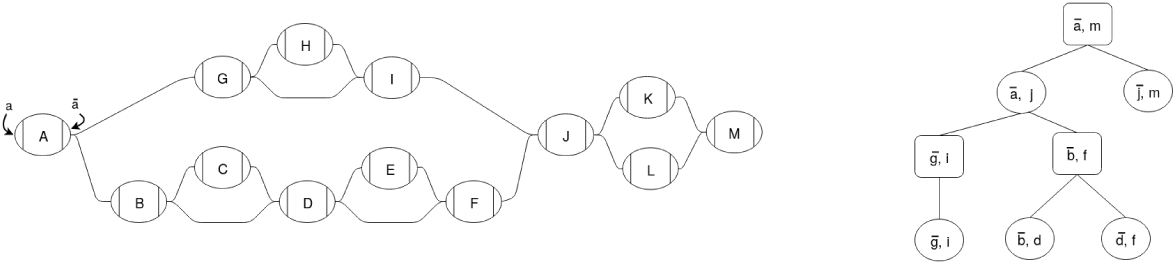
Example sequence graph (left) and its snarl tree (right). Chains in the sequence graph are represented as rectangular nodes in the snarl tree and snarls are represented as elliptical nodes.

Some applications use a specific articulation of a sequence graph called a variation graph. A variation graph contains a set of embedded paths through the graph, frequently including the path corresponding to the linear reference.

### 2.2 Snarl Decomposition

In previous work, we proposed a decomposition for sequence graphs that describes their common topological features [12]. A simple variant in a graph, such as an indel or SNP, will typically be represented as one or two nodes that represent the different alleles, flanked by two nodes representing conserved sequences. The subgraph between the two flanking nodes is called a snarl. A snarl is defined by a pair of node sides, (*x, y*) that delimit a subgraph between them. The nodes *X* and *Y* are called the boundary nodes of the snarl. Two node sides define a snarl if they are (1) separable: splitting the boundary nodes into their two node sides disconnects the snarl from the rest of the graph, and (2) minimal: there is no node *A* in the snarl such that (*x, a*) or (*ā, y*) are separable. (Note that unary snarls, where *x* = *y*, are permitted only at “dead ends” where ordinary snarls are not possible.) We refer to the snarl defined by node sides *x* and *y* as (*x, y*). In Figure 1, 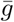 and *i* define a snarl, 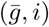, that contains node *H*.

In sequence graphs, snarls often occur contiguously with a shared boundary node between them; a sequence of contiguous snarls is called a chain. In Figure 1, the snarls 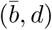 and 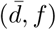 comprise a chain between 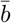 and *f*, which we refer to as 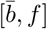. A trivial chain is one that contains only one snarl; in Figure 1, snarl 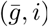 is part of a trivial chain, chain 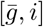.

Snarls and chains can be nested within other snarls. A snarl (*x, y*) contains another snarl (*a, b*) if all nodes in (*a, b*) are contained in the subgraph of (*x, y*). In Figure 1, the snarl (*ā, j*) contains snarls 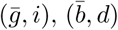, and 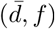. A snarl contains a chain if each of the chain’s snarls are in the subgraph of the containing snarl.

The nesting relationships of snarls and chains in a sequence graph is described by its snarl tree (Figure 1). Each snarl or chain is represented in the snarl tree as a node. Since every snarl belongs to a (possibly trivial) chain, snarl trees have alternating levels of snarls and chains with a chain at the root of the tree. A snarl is the child of a chain if it is a component of the chain. A chain [*a, b*] is a child of (*x, y*) if (*x, y*) contains [*a, b*] and there are no snarls contained in (*x, y*) that also contain [*a, b*].

Nodes, snarls, and chains are all two-ended structures that are connected to the rest of the graph by two node sides. (Unary snarls can occur only at dead ends in the graph; their “other sides” are disconnected.) It is sometimes convenient to refer to a topological feature only by this shared property, and to be opaque about which topological feature it actually is. In these cases, we will refer to the node, snarl, or chain generically as a *structure*. As with nodes of the sequence graph itself, structures are assigned an arbitrary orientation but we will assume that they are oriented left to right and refer to the left and right sides of structures as *struct* and 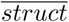 respectively. Because of their shared two-ended property, structures can all be treated as single nodes in their parents. The netgraph of a snarl is a view of the snarl where each its child chains is replaced by a node.

### 2.3 Prior Research

#### Distance in graphs

Calculating distance in a graph is an extremely well studied topic. Many graph distance algorithms improve upon classical algorithms, such as Dijkstra’s algorithm [4] and A* [7], by storing precomputed data in indexes. These methods index the identities of important edges [11, 9] or distances between selected nodes [3, 14, 5, 2] then use the indexed information to speed up distance calculations. Index-based algorithms must make a tradeoff between the size of the index and the speed of the distance query.

#### Distance in sequence graphs

Some sequence graph mapping algorithms use clustering steps based on different estimations of distance [18, 6]. In `vg`, distance is approximated from the haplotype paths. This path-based method finds the distance between two positions on a shared haplotype path. If there is no shared path, then the algorithm traverses the graph from each positions until it finds a path reachable from both positions and returns the sum of the distances to the path and distance between the positions in the path.

Some research has been done on finding solutions for more specific distance queries in sequence graphs. PairG [8] is a method for determining the validity of independent mappings of reads in a pair by deciding whether there is a path between the mappings whose distance is within a given range. This algorithm uses an index to determine if there is a valid path between two vertices in a single O(1) lookup. Although this is an efficient solution for this particular problem, it cannot be used for dynamic distance queries since it returns a boolean value of whether two nodes are reachable within a range of distances, which is defined at index construction time.

## 3 Minimum Distance

Our minimum distance algorithm finds the minimum oriented traversal distance between two positions on a sequence graph. A position consists of a node, offset in the sequence, and orientation. The oriented distance must originate from a path that starts traversing the first position in its given orientation and ends at the second position in its given orientation.

Our algorithm uses the snarl decomposition of sequence graphs to guide the calculation. Because structures are connected to the rest of the graph by their boundary nodes, any path from a node inside a structure to any node not in that structure must pass through the structure’s boundary nodes. Similarly, any path between boundary nodes of snarls in a chain must pass through the boundary nodes of every snarl that occurs between them in the chain. Because of this property, we can break up the minimum distance calculation into minimum distances from node and chain boundaries to the boundaries of their parent snarl, from snarl boundaries to their parent chain boundaries, and the distance between sibling structures in their parent structure (Figure 2). We refer to this property of minimum distance calculation in structures as the split distance property.

**Fig. 2.**
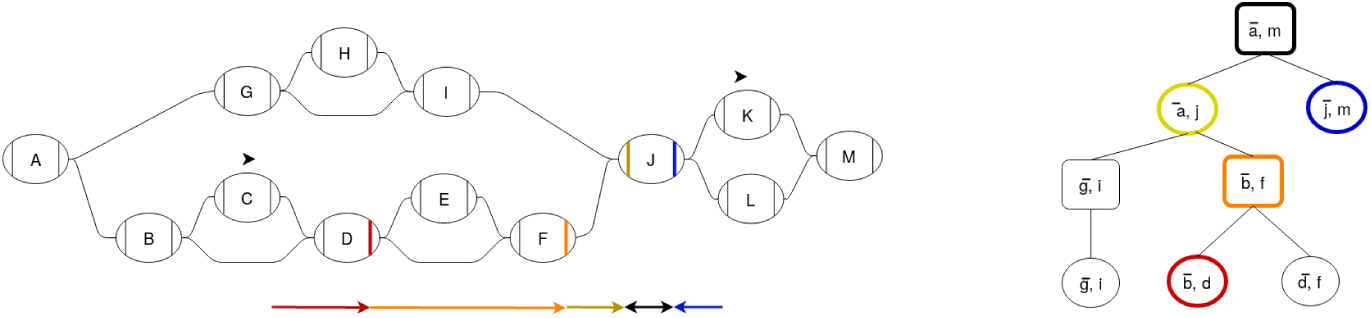
The minimum distance calculation from a position on *C* to a position on *K* can be broken up into the distances from each position to the ends of each of its ancestor structures in the snarl tree. Each colored arrow in the graph represents a distance query from a structure to a boundary node of its parent. The snarl tree node that each query occurs in is outlined with the same color. At the common ancestor of the positions, chain [*ā, m*], the distance is calculated between two of the chain’s children, (*ā, j*) and 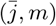.

### 3.1 Minimum Distance Index

We designed our minimum distance index to support distance queries between child structures in snarls and between boundary nodes of snarls in chains in constant time. The overall minimum distance index consists of a snarl index for each snarl and a chain index for each chain in the graph.

#### Snarl Index

For each snarl, the index stores the minimum distances between every pair of child structures. A distance query within a snarl is a simple constant time lookup of the distance.

#### Chain Index

For each chain, the index stores three arrays, each with one entry for each boundary node of the snarls in the chain. A prefix sum array contains the minimum distance from the start of the chain to the boundary nodes of each of the snarls that comprise the chain. Since paths can reverse direction in the chain (Figure 3), the index also stores the minimum distance to leave the boundary node, change direction in the chain, and return to the same node side. These loop distances are stored in two arrays, one for traversing the chain forward and one for traversing it backward. These three arrays are sufficient to find the minimum distance between any two node sides in the chain in constant time (Figure 3).

**Fig. 3.**
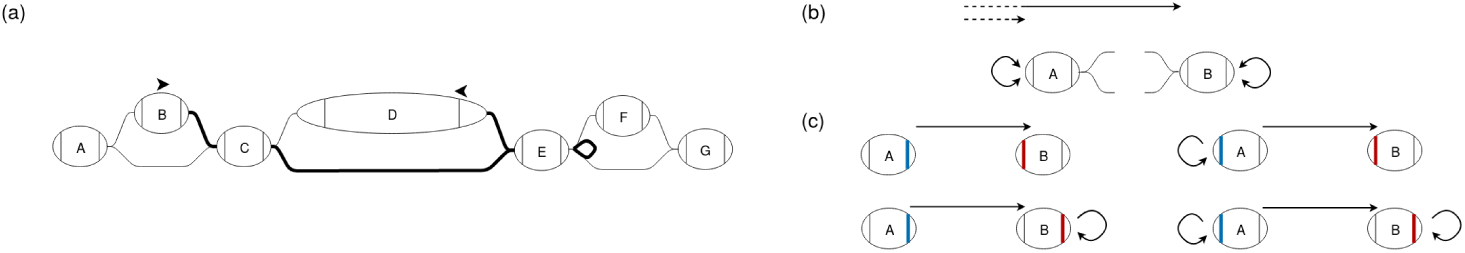
(a) The shortest path between two nodes in a chain can sometimes reverse direction in the chain. The edges on the shortest path between the positions on *B* and *D* are bolded. (b) *A* and *B* are boundary nodes of snarls in a chain. Distances stored in the chain index are shown in black. For each boundary node in the chain, the chain index stores the minimum distance from the start of the chain to that node as well as the loop distances for a forward and backward traversal. These loop distances are the minimum distance to leave a node, reverse direction in the chain, and return to the same node side. (c) There are four possible minimum-distance paths between two nodes, connecting either node side of the two nodes. The lengths of these paths can be found using the distances stored in the chain index.

Chains that are not top-level chains cannot form a closed cycle so any path that traverses a chain’s boundary node going out of the chain must leave the chain. Therefore any connectivity between the boundaries of the chain will be captured by the snarl index of the chain’s parent. The top-level chain may form a closed cycle where the start and end boundary nodes are the same node (Figure 4). In this case, the shortest path may remain within the chain, but it may also leave the chain and re-enter it from the other side. In Figure 4, the minimum distance from *ā* to *d* could be 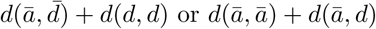.

**Fig. 4.**
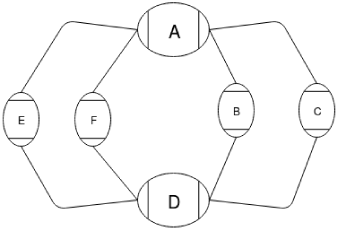
A cyclic chain containing two snarls, 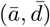 and (*a, d*)

#### Index Construction

The minimum distance index is constructed in a post-order traversal of the snarl tree. For each snarl, the construction algorithm does a Dijkstra traversal starting from each child structure, using the child’s index to find the distance to traverse child snarls or chains. For each chain, the construction algorithm traverses through each snarl in the chain and uses the snarl’s index to find each of the relevant distances for the chain index.

#### Index Size

Naively, a minimum distance index could store the minimum distance between every node in the graph. A distance calculation would be a constant time lookup but the index size would be quadratic in the number of nodes in the graph. For each snarl in the graph, our index stores the distance between every pair of structures in the net graph. For each chain, it stores three arrays, each the length of the chain. In a graph with a set of snarls *S* and chains *C*, our index will take 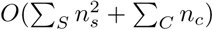 space where *n*_*s*_ is the number of structures in the netgraph of snarl *s* and *n*_*c*_ is the number of snarls in chain *c*.

### 3.2 Algorithm

The first step of our minimum distance algorithm is to find the least common ancestor structure in the snarl tree that contains both positions. We do this by traversing up the snarl tree from each position and finding the first common structure. This traversal is *O*(*d*) where *d* is the depth of the snarl tree.

Next, the algorithm finds the distance from each position to the ends of the child of the least common ancestor. Starting at a node containing a position, we find the distances to the ends of the node. Since we are finding the oriented distance, one of these distances is infinite. The algorithm then traverses up the snarl tree to the least common ancestor and at each structure, finds the minimum distances to the ends of the structure. Because of the split distance property, this distance can be found by adding the distances to the ends of the child, found in the previous step in the traversal, to the distances from the child to the boundary nodes of the structure, found using the minimum distance index (Figure 5). Since this requires only four constant-time queries to the minimum distance index, each step in the traversal is constant time and the overall traversal is *O*(*d*).

**Fig. 5.**
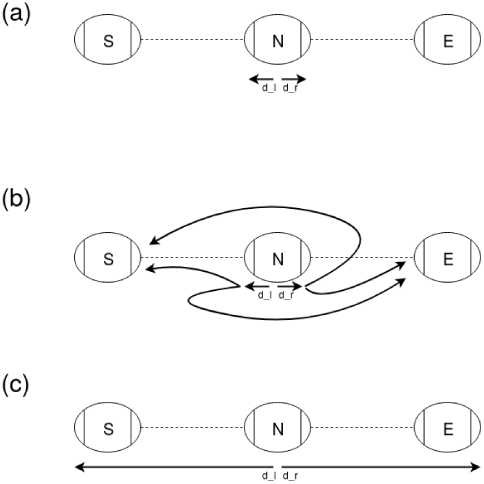
(a) *S* and *E* are the boundary nodes of a structure that contains a child structure *N*. *d*_*l*_ and *d*_*r*_ are the minimum distances from some object in *N* to the ends of *N*. (b) The minimum distances from each end of *N* to 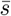 and *e* are found using the minimum distance index. (c) By adding the appropriate distances and taking the minimums, we can get the minimum distances to *s* and *ē*.

At this point in the algorithm, we know the minimum distance from each position to its ancestor structure that is a child of the common ancestor. By composing these distances with the distances between the two structures, the algorithm finds possible distances between the two positions in the common ancestor structure. The algorithm continues to traverse the snarl tree up to the root and finds a minimum distance between the positions at each structure, checking for paths that leave the lowest common ancestor. This traversal is also *O*(*d*). The minimum distance algorithm is done in three *O*(*d*) traversals of the snarl tree, so the algorithm is *O*(*d*). In real sequence graphs, the maximum depth of the snarl trees is very small so in practice our algorithm is expected to be *O*(1).

**Table 1.** Primitive functions for the minimum distance algorithm.

| Function | Description | Complexity |
| --- | --- | --- |
| `distToEndsOfParent(struct, dist_left, dist_right)` | Given the distances from a position in a structure *struct* to the ends of *struct*, find the distance to the ends of the parent (Figure 5) | $O(1)$<br>using the distance index |
| `distWithinStructure(struct, child_1, child_2, dist1_l, dist1_r, dist2_l, dist2_r)` | Given two children of a structure and distances from positions to the boundaries of the children find the minimum distance between the positions in *struct* | $O(1)$<br>using the distance index |

#### Algorithm 1: distToAncestor(*position, ancestor*) Given a position and ancestor structure, return the minimum distance from the position to both sides of a child of the ancestor and the child

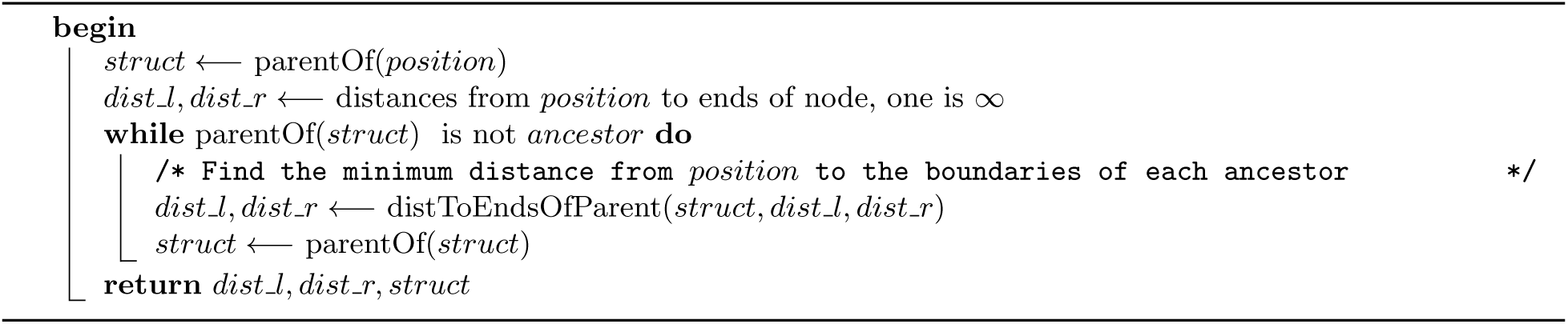

## 4 Clustering

Seed-and-extend algorithms sometimes cluster seed alignments by their location in the graph to find which might belong to the same mapping. Using our minimum distance index, we developed an algorithm to cluster positions based on the minimum distance between them in the graph.

### 4.1 Problem

We will cluster seeds by partitioning them based on the minimum distance between their positions in a sequence graph. To define a cluster, we consider a graph where each seed is a node and two seeds are connected if the minimum distance between their positions is less than a given distance limit. In this graph, each connected component is a cluster.

### 4.2 Algorithm

Our clustering algorithm starts with each position in a separate cluster then progressively agglomerates the clusters (Figure 6). The algorithm proceeds in a post-order traversal of the snarl tree and, at each structure, produces clusters of all positions contained in that structure (Algorithm 5). After iterating over a structure, clusters are also annotated with two “boundary distances”: the shortest distance from any of its positions to the boundary nodes of the structure. At every iteration, each cluster can be unambiguously identified with a structure and so the boundary distances are always measured to the structure the cluster is on.

**Fig. 6.**
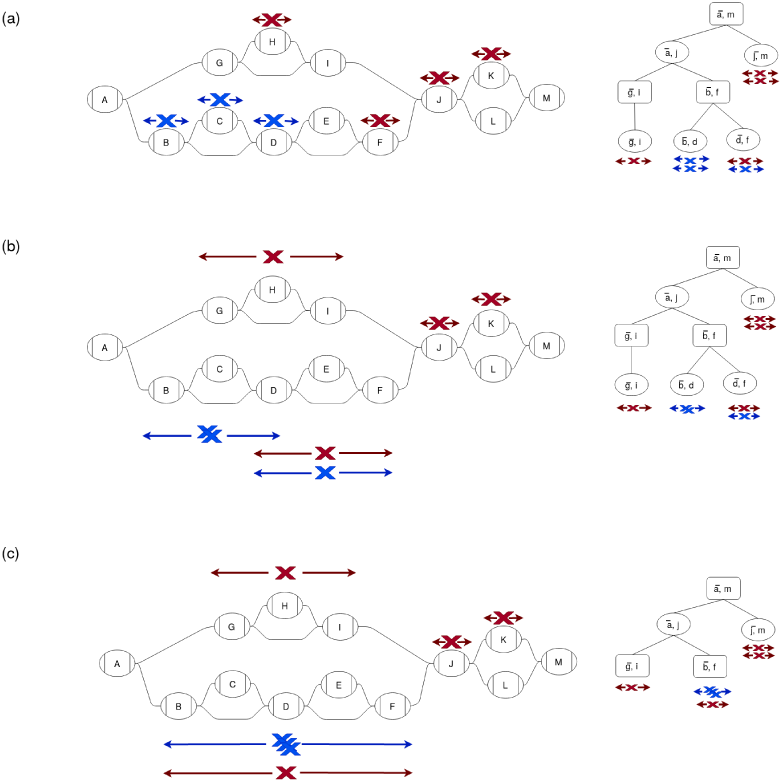
Clustering of positions (Xs) is done by traversing up the snarl tree and progressively agglomerating clusters. Positions are colored by the final clusters. (a) Each position starts out in a separate cluster on a node. Each cluster is annotated with its boundary distances: the minimum distances from its positions to the ends of the structure it is on. (b) For each snarl on the lowest level of the snarl tree, the clusters on the snarl’s children are agglomerated into new clusters on the snarl. The boundary distances are extended to the ends of the snarl. (c) For each chain on the next level of the snarl tree, the clusters on the chain’s snarls are agglomerated and the boundary distances are updated to reach the ends of the chain. This process is repeated on each level of the snarl tree up to the root.

The method of agglomerating clusters and computing boundary distances vary according to the type of structure. For nodes, the algorithm creates a sorted array of the positions contained in it and splits the array into separate clusters when the distance between successive positions is large enough. For each new cluster, the boundary distances are computed from the positions’ offsets.

For structures that are snarls or chains, clusters are created from the clusters on their children (Algorithm 3, Algorithm 4). Clusters associated with child structures are compared and if the distance between any pair of their positions is smaller than the distance limit, they are combined. Within a structure, distances to clusters that are associated with child structures can be calculated using the split distance property as in the minimum distance algorithm. According to this property, the minimum distance can be split into the cluster’s boundary distance and the distance to one of the boundary nodes, which is found using the index. For snarls, all pairs of clusters are compared to each other. For chains, clusters are combined in the order they occur the chain, so each cluster is compared to agglomerated clusters that preceded it in the chain. Finally, for each of the resulting clusters, we compute the boundary distances for the current structure, once again using the boundary distances of the children and the index.

#### Algorithm 2: minDistance(*position_*1, *position_*2) Return the minimum distance from *position_*1 to *position_*2, ∞ if no path between them exists

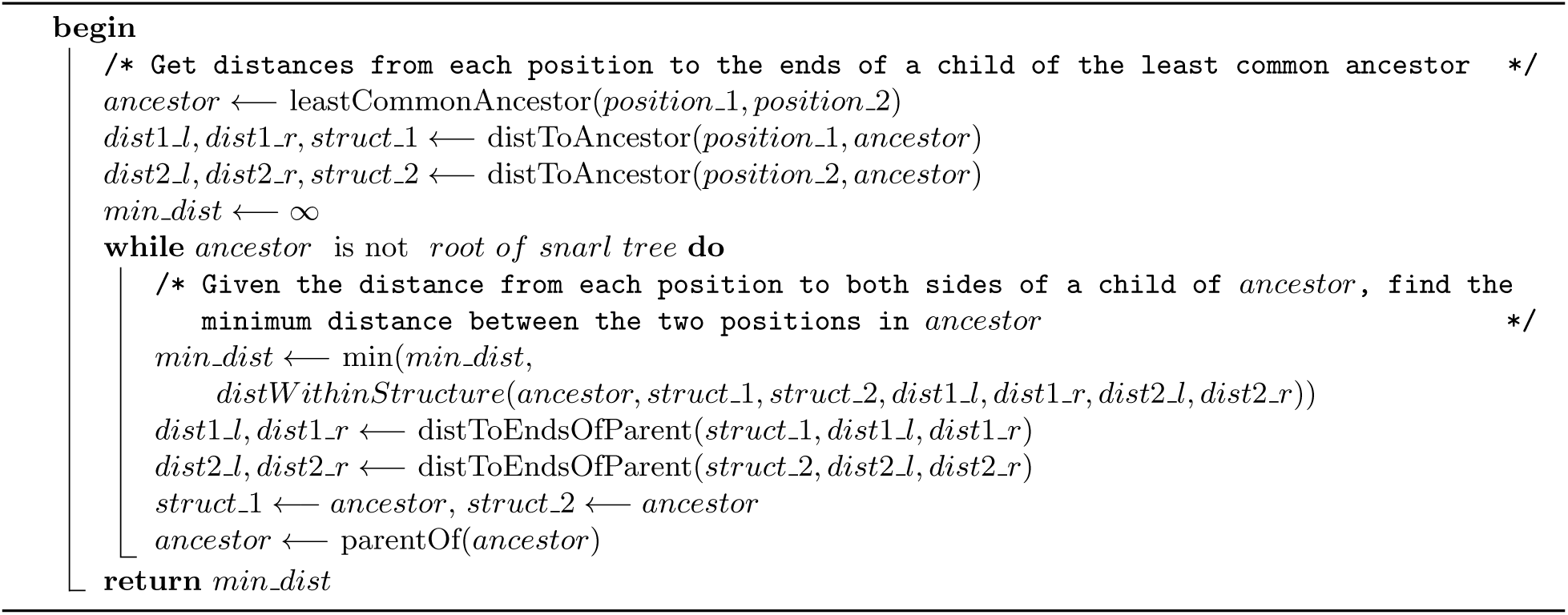

In the worst case, every position would belong to a separate cluster and at every level of the snarl tree, ever cluster would be compared to every other cluster. This would be *O*(*dn*^2^) where *d* is the depth of the snarl tree and *n* is the number of seeds, so in the worst case our clustering algorithm is no better than the naive algorithm of comparing every pair of positions with our minimum distance algorithm. In practice however, seeds that came from the same alignment would be near each other on the graph form clusters together, significantly reducing the number of distance comparisons that would be made (see results below).

## 5 Methods and Results

Our algorithms are implemented as part of the `vg` toolkit. We conducted experiments on two different graphs: a human genome variation graph and a graph with simulated structural variants. The human genome variation graph was constructed from GRCh37 and the variants from the 1000 Genomes Project. The structural variant graph was simulated with 10bp-1kbp insertions and deletions every 500bp.

The human genome variation graph graph had 306, 009, 792 nodes, 396, 177, 818 edges, and 3, 180, 963, 531 bps of sequence. The snarl tree for this graph had a maximum depth of three snarls with 139, 418, 023 snarls and 11, 941 chains. The minimum distance index for the graph was 12.2 GB on disk and 17.7 GB in memory.

To assess the run time of our minimum distance algorithm, we calculated distances between positions on the whole genome graph and compared the run time of our algorithm to `vg`’s path-based algorithm and Dijkstra’s algorithm (Figure 7). We chose random pairs of positions in two ways. The first method sampled positions uniformly at random throughout the graph. The second method first followed a random walk of 148 bp through the graph and then sampled two positions uniformly at random from this random walk. This approach was intended to approximate the case of seeds from a next-generation sequencing read. On average, our minimum distance algorithm is the fastest of the three algorithms for both sets of positions. In addition, all three algorithms’ performance degraded when the positions could be sampled arbitrarily far apart in the graph, but our minimum distance algorithm’s performance degraded the least.

**Fig. 7.**
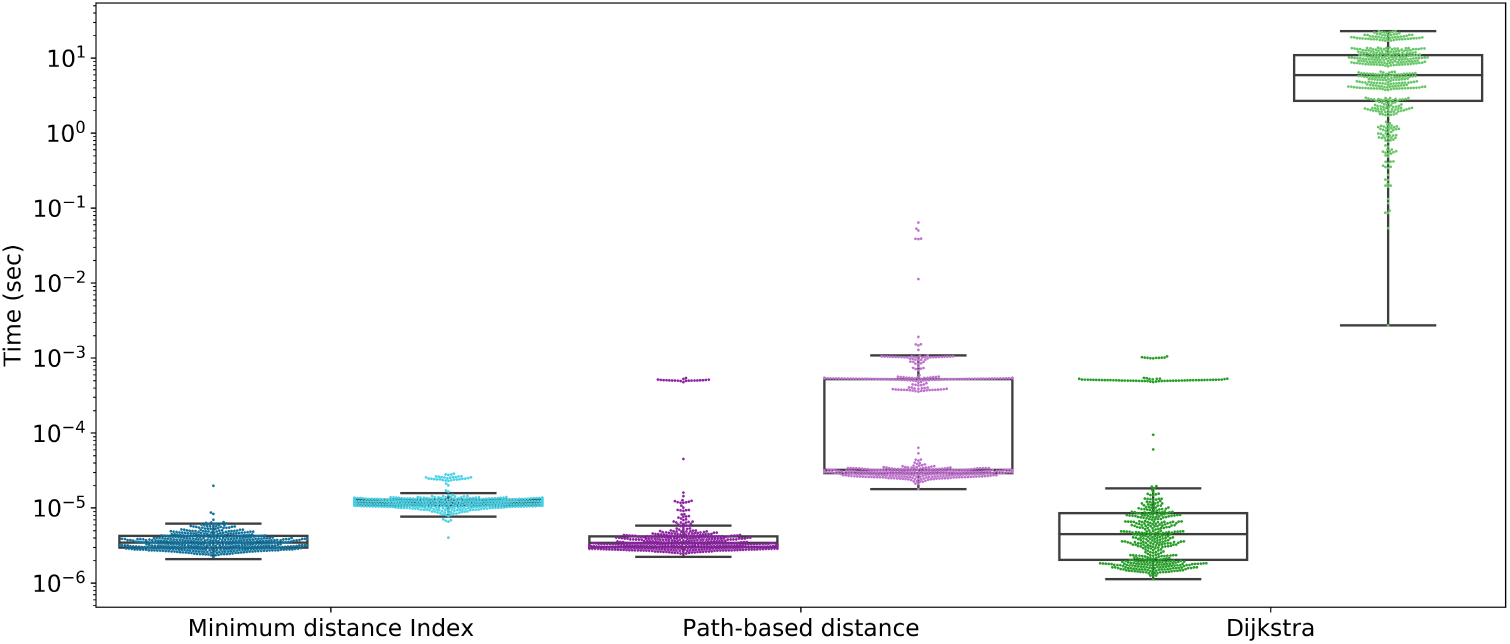
Run times for distance algorithms. Random pairs of positions were chosen from either within a read-length random walk (dark colors) or randomly from the graph (light colors).

We used the structural variant graph to assess whether the minimum distance is a useful measure of distance for read mapping. To do so, we again used read-length random walks to select pairs of positions. Further, we filtered random walks down to those that overlapped a structural variant breakpoint. We then calculated the distances between pairs of positions using our minimum distance algorithm and the path-based approximation and compared these distances to the actual distances in the random walk, which we take as an approximation of the true distance on a sequencing read. Overall, the minimum distance was a much better estimate of distance along the random walk than the path-based distance approximation (Figure 8).

**Fig. 8.**
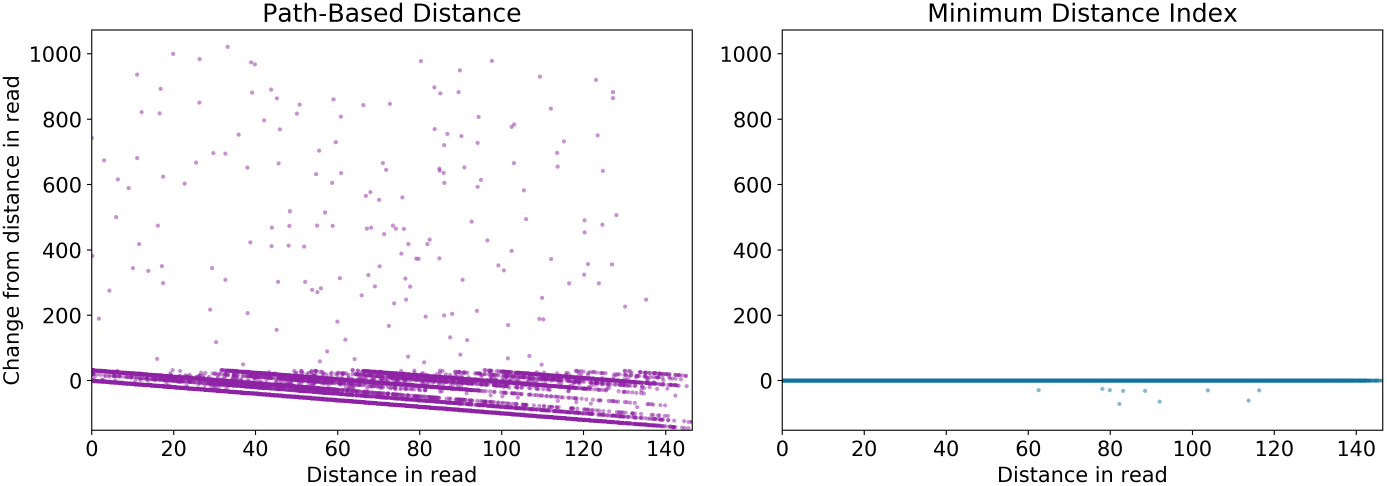
Distance calculations on a graph with simulated structural variants. Read-length random walks were simulated near the junctions of structural variants. The distance between two random positions along each walk was calculated using our minimum distance algorithm (top) and the path-based method (bottom) and compared to the actual distance in the walk.

**Fig. 9.**
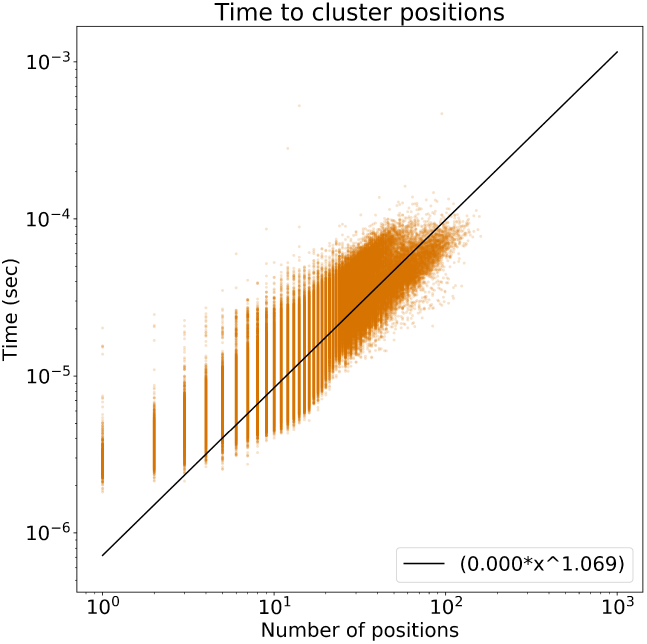
Run time growth of our clustering algorithm. The regression line suggests that the run time of our algorithm is nearly linear in the number of positions.

For our clustering algorithm, we wanted to estimate the run time of the algorithm in the context of read mapping. We simulated 148bp reads from AshkenazimTrio HG002_NA24385_son from the Genome in a Bottle Consortium [19]. For each read, we sampled 15-mer matches from the read and found their positions in the human genome variation graph using a *k*-mer lookup table. We then apply the clustering algorithm to the positions of these *k*-mers. The regression line of the log-log plot of run times suggests the run time of our algorithm is linear in the number of positions in practice, despite the quadratic worse-case bound.

### Algorithm 3: clusterSnarl(*snarl, child_to_clusters, distance_limit*) Given a snarl and map from children of the snarl to their clusters, get clusters of the snarl

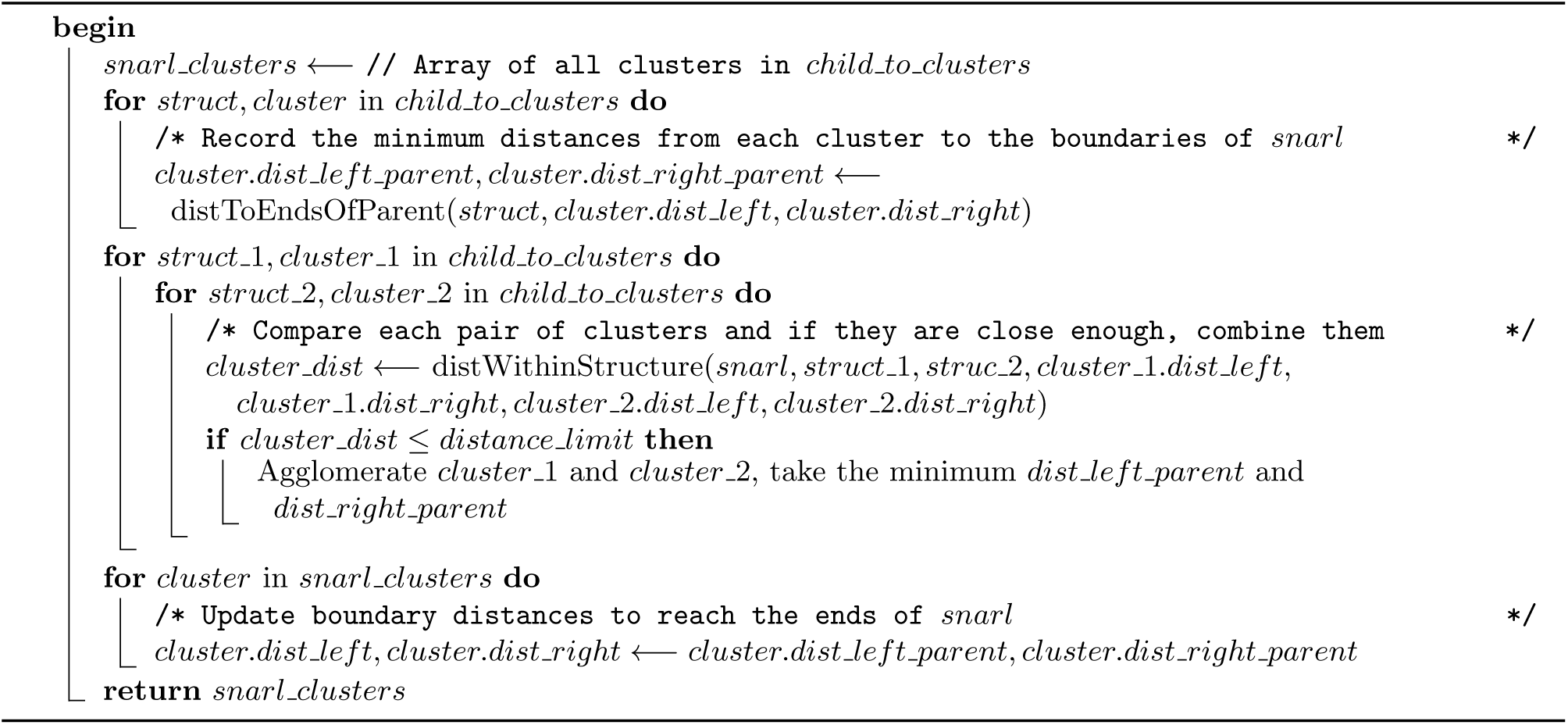

## 6 Conclusion

We developed a minimum distance algorithm with run time that is linear in the depth of the snarl tree. In practice, the algorithm exploits the observation that real-world genome graphs have an excess of small, local variations and relatively fewer variations that connect disparate parts of the graph. The result is that real genome graphs have a shallow snarl tree, making the calculations fast and close to constant time in practice. The distance we return is also more strictly defined than the previous implementation of distance in `vg` and is faster than other distance algorithms on queries of arbitrary distance. Our minimum distance algorithm will also work with any sequence graph, whereas the preexisting `vg` distance algorithm required pre-specified paths. We also developed a clustering algorithm for clustering positions on the graph based on the minimum distances between them. We are developing fast mapping algorithms that use this clustering algorithm.

### Algorithm 4: clusterChain(*chain, child_to_clusters, distance_limit*) Given a chain and a map from each snarl in the chain to its clusters, get clusters of the chain

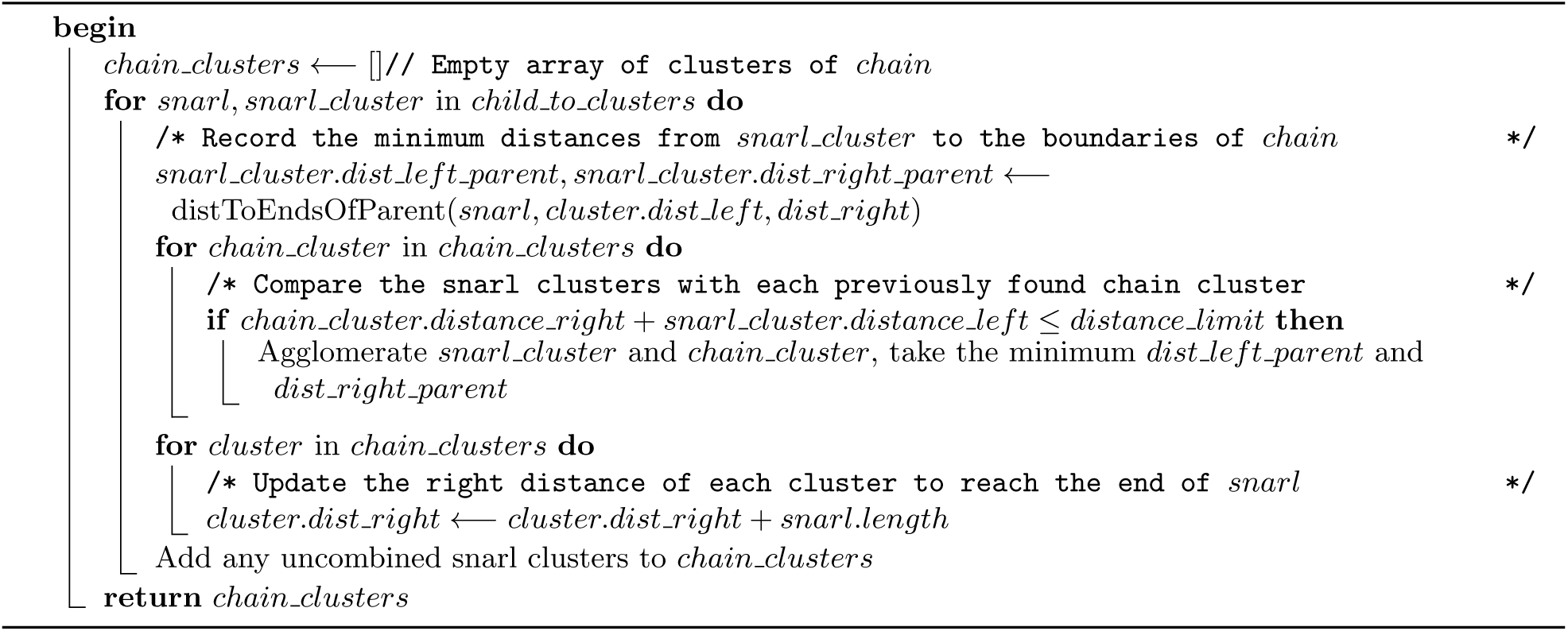

### Algorithm 5: cluster(*snarl_tree, positions, distance_limit*) Cluster positions based on the distance limit

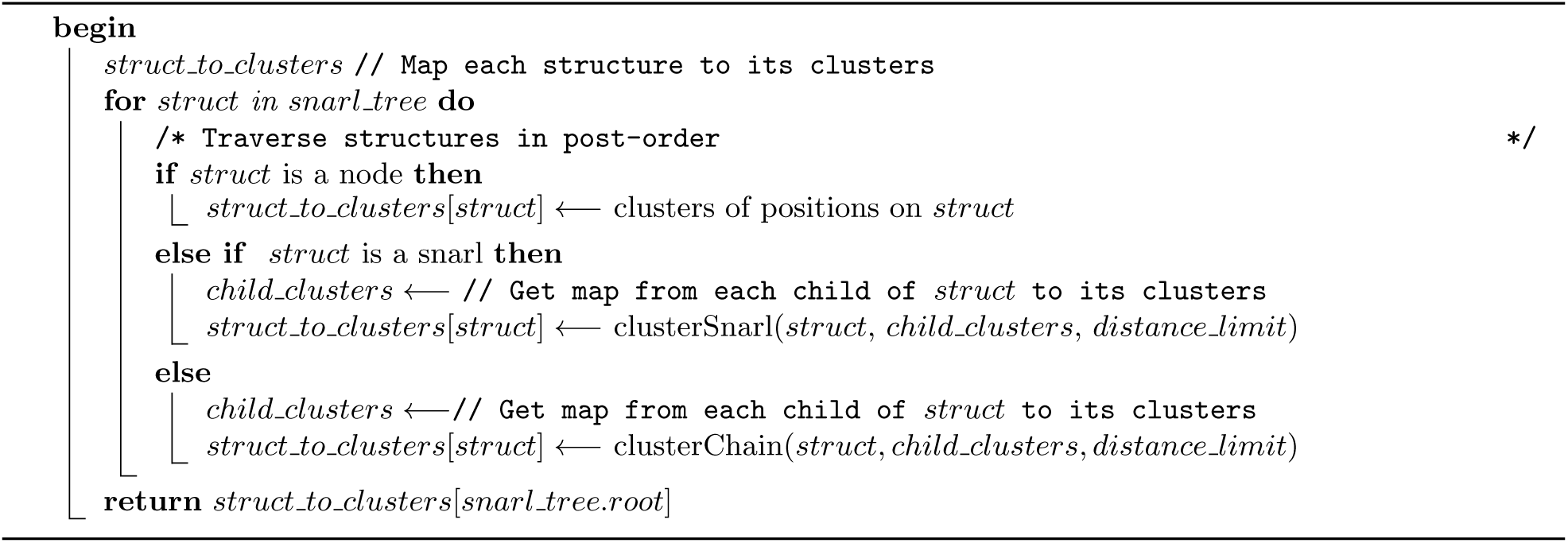

